# Efficient and accurate detection of splice junctions from RNAseq with Portcullis

**DOI:** 10.1101/217620

**Authors:** Daniel Mapleson, Luca Venturini, Gemy Kaithakottil, David Swarbreck

## Abstract

Next generation sequencing (NGS) technologies enable rapid and cheap genome-wide transcriptome analysis, providing vital information about gene structure, transcript expression and alternative splicing. Key to this is the the accurate identification of exon-exon junctions from RNA sequenced (RNA-seq) reads. A number of RNA-seq aligners capable of splitting reads across these splice junctions (SJs) have been developed, however, it has been shown that while they correctly identify most genuine SJs available in a given sample, they also often produce large numbers of incorrect SJs. Herein we describe the extent of this problem using popular RNA-seq mapping tools, and present a new method, called Portcullis, to rapidly filter false SJs junctions from spliced alignments produced by any RNA-seq mapper capable of creating SAM/BAM files. We show that Portcullis distinguishes between genuine and false positive junctions to a high-degree of accuracy across different species, samples, expression levels, error profiles and read lengths. Portcullis makes efficient use of memory and threading and, to our knowledge, is currently the only SJ prediction tool that reliably scales for use with large RNAseq datasets and large highly fragmented genomes, whilst delivering highly accurate SJs.

**Availability:** Portcullis is available under the GPLv3 license at: http://maplesond.github.io/portcullis/

## 1 INTRODUCTION

Alternative splicing (AS) is a regulated process in Eukaryotic species that occurs during gene expression enabling a single gene to code for multiple proteins through inclusion or exclusion of exons in the transcribed mRNA. Key to defining the complexity of alternative splicing within a gene is the identification of splice junctions, which occur at exon-exon boundaries and are typically characterised in pairs representing both the donor site (5’ intron boundary to 3’ upstream exon boundary) and acceptor site (3’ intron boundary to 5’ downstream exon boundary). A recent study has given us a comprehensive view into alternative splicing in Humans (Nellore *et al.*, 2016), although annotations of other model species are known to be incomplete (Robert *et al.*, 2014). This lack of completeness reduces the accuracy and usefulness of many downstream gene and transcript level tasks, such as differential expression analysis (Soneson and Delorenzi, 2013) and alternative splicing analysis (Christinat *et al.*, 2016).

RNAseq has become the standard method to detect, quantify, compare and contrast splice isoforms across different biological contexts (Conesa *et al.*, 2016). Furthermore, as Next Generation Sequencing (NGS) technologies have matured, RNA-seq is becoming increasingly quick, reliable and cost-effective (Goodwin *et al.*, 2016). Splice junctions derived from RNAseq studies are primarily detected via RNA-seq mappers, which split reads derived from transcripts across introns. However, a survey of many RNA-seq mapping tools highlighted that accurate detection of splice junctions is an outstanding challenge (Engström *et al.*, 2013). We show in this manuscript that this issue persists with the latest versions of many popular RNAseq mappers. The lack of accuracy is due to various factors, such as short read lengths increasing mapping ambiguity, sequencing errors triggering misaligned split reads, and is exacerbated in deeply covered datasets where the likelihood of generating distinct invalid splice sites increases.

There are a number of strategies for reducing the number of distinct spurious SJs found in mapped reads. One method involves counting the split reads supporting each distinct junction and filtering those under a certain threshold (Nellore *et al.*, 2016). Other methods utilise the length of split read overhangs across each junction from the set of supporting split reads (Wang *et al.*, 2010b), or calculating the amount of evidence supporting their start sites (Wang *et al.*, 2010a). Studying the genome around the splice junction has also proved effective (Huang *et al.*, 2011; Li *et al.*, 2013; Gatto *et al.*, 2014). To reduce the propagation of invalid junctions into downstream tasks, many RNAseq mappers offer the ability to use a set of high-confidence junctions to guide realignment of the reads.

In practice, the computational demands of collecting a set of accurate junctions rapidly can be problematic, often leading to compromised accuracy. Accurate methods either require long runtimes, high-memory usage or are inflexible and difficult to use. We address these issues with our tool, Portcullis, which is the only method we are aware of, for rapidly and accurately filtering invalid SJs from BAM alignments produced by any RNA-seq mapper. Portcullis competes with or outperforms the best methods currently available in terms of accuracy, speed and memory efficiency, over a wide range of scenarios. Portcullis also offers rich junction analysis and quantification capabilities, as well as a supplementary toolkit, called Junctools, for performing tasks such as junction file format conversions, junction set comparisons and various filtering options.

## 2 RESULTS

### 2.1 Junction detection performance

It has previously been observed that short-read RNA-seq mapping tools often produce large numbers of false positive junctions (Engström *et al.*, 2013). To gauge the extent of this problem with a more recent set of popular mapping tools, we generated several sets of simulated reads with varying read length and depth from 3 different species transcriptomes (see section 5.1). We then extracted a distinct set of SJs from split reads produced by the mappers and compared them to the set of true junctions for each corresponding simulated dataset. We use the recall, precision and *F*_1_ (F-measure) to assess the performance of each mapper (see section 5.2 for a description of how these are calculated). Figure 1(a) shows the performance of STAR v2.5.2a for four 201bp paired-end read datasets of the Human transcriptome, each with varying depth. Figure 1(b) shows the effect of varying read length, with each dataset having a depth of ~30 billion base pairs. The plots show that a longer read length improves both recall and precision, however increased depth, while marginally improving recall, decreases precision significantly, pulling *F*_1_ down with it. This decrease in precision is in part attributable to reads containing sequencing errors triggering misalignments of split reads, leading to new, invalid, junctions being predicted (Wang *et al.*, 2010a). Supplementary Information (SI) section 4 shows the same trends hold true for several other popular RNA-seq mappers such as: TopHat2 v2.1.0(Kim *et al.*, 2013), GSNAP v20160923(Wu and Nacu, 2010) and HISAT v2.0.5(Kim *et al.*, 2015). In addition, figure 2 highlights that the effect is not limited to human, a species with complex splicing behaviour, but is also visible in species with significantly less splicing events such as *Arabidopsis* and *Drosophila*.

**Fig. 1:**
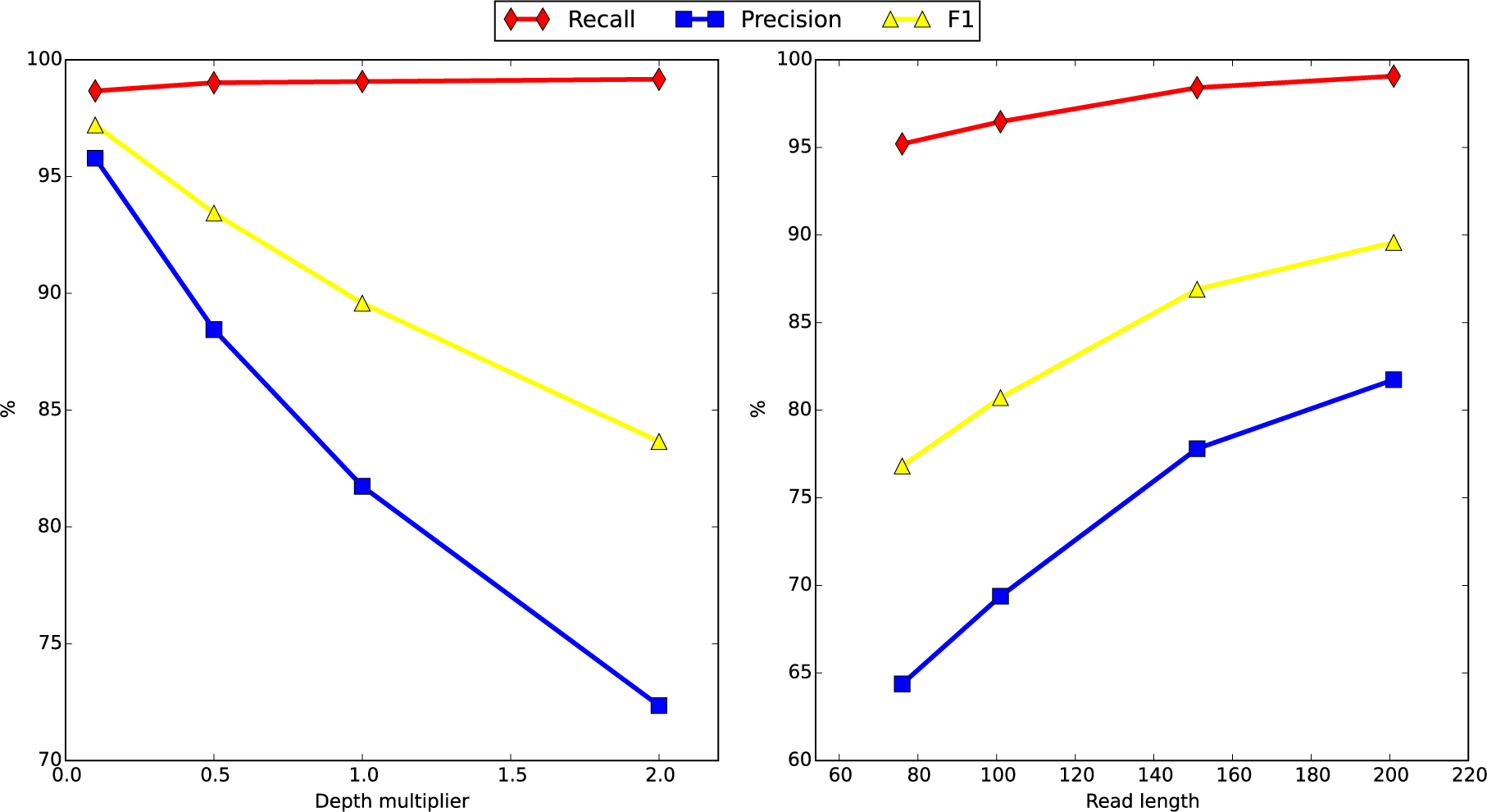
Splice junction accuracy of STAR v2.5.2a across variations of our simulated human dataset. (a) - left - shows the effect of varying dataset size, with all datasets containing 201bp reads. 1.0X depth multiplier represents a dataset of ~78 million read pairs. (b) - right - shows the effect of varying read length with all datasets containing ~30 billion base pairs.

**Fig. 2:**
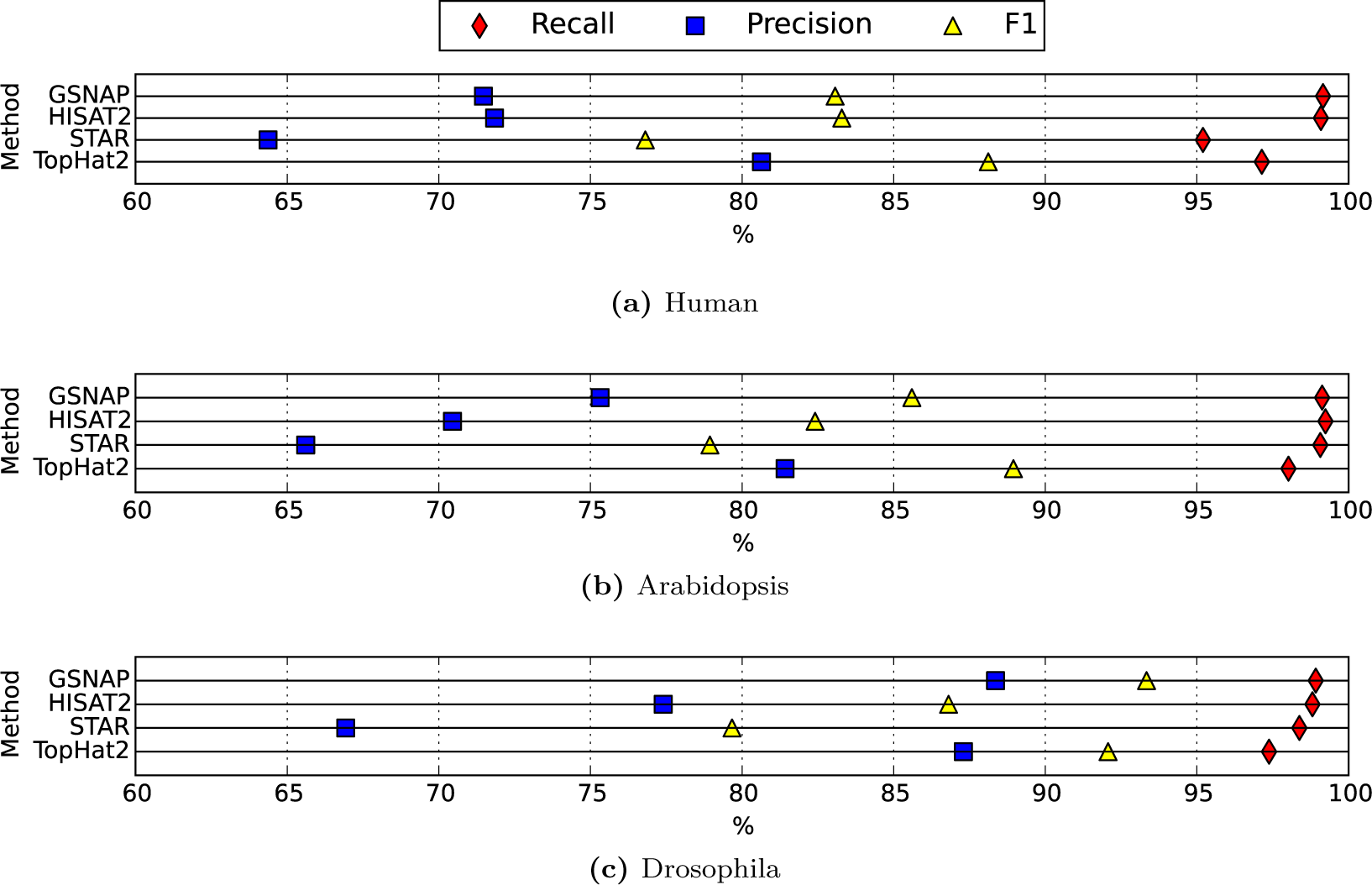
Splice junction detection performance across mappers for 76bp simulated paired reads. The Human dataset (a) contains 421, 020, 756 reads across 19, 853 transcripts. The *Arabidopsis* dataset (b) contains 148, 207, 902 reads across 19, 723 transcripts. The *Drosophila* dataset (c) contains 202, 246, 654 reads across 9, 376 transcripts.

We also observed that, while all of these mappers recall a good fraction of genuine junctions, the mappers are not producing the same sets of false positives. The Venn diagram in figure 3 shows the agreement between junctions found across Tophat2, GSNAP, STAR, and HISAT2 mappers on our 76bp simulated human dataset. Each mapper retrieves at least 93.6% of true junctions, although there is little agreement for the remaining junctions, indicating that these mappers often make different types of mistakes. SI figure 4 shows the effect of different depth and read length but essentially the same trends hold. These observations indicate that a high-confidence set of splice junctions can be built by requiring a degree of concordance between multiple mappers, which we demonstrate in SI figure 5. As the expected number of mappers required to agree increases, there is a corresponding increase in precision at the expense of recall. We note that in the cases when 2, 3 or 4 mappers are required to agree, the *F*_1_ exceeds that of any of the individual mapping tools. However, this approach has several disadvantages. First, it assumes that the aligners find different false positives. While this assumption is true for our selection of mappers, it may not always be true as different mappers (or mapper versions) may have different characteristics. Second, a single poorly performing mapper will reduce the performance of the system as a whole. Third, it is not possible to know *a priori* how much agreement is required to get optimal results. Finally, this approach is not particularly computationally efficient as it requires running multiple tools for a single dataset.

**Fig. 3:**
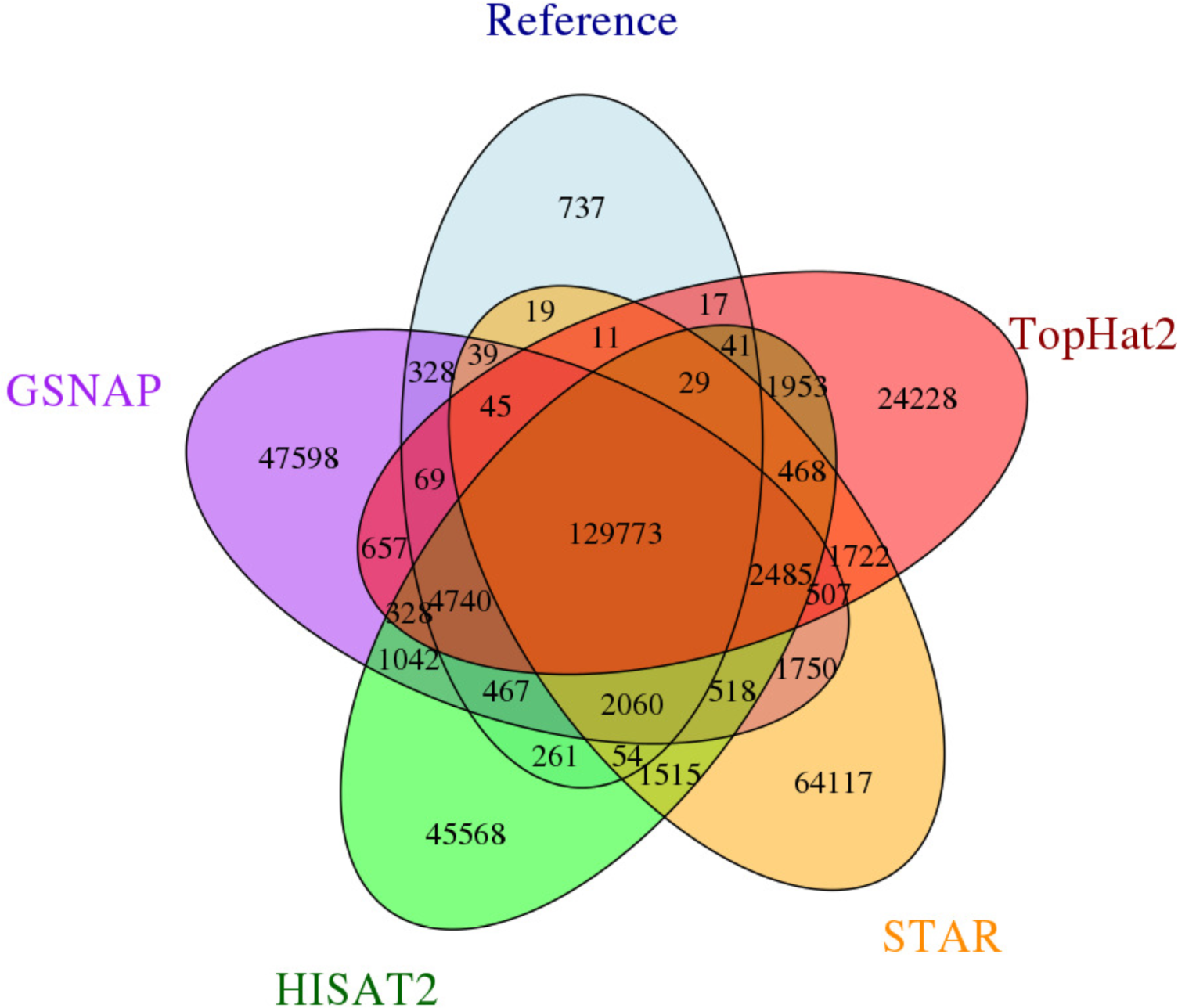
A five way Venn diagram showing levels of agreement between mapping tools and the human junction truth set with 76bp simulated reads.

**Fig. 4:**
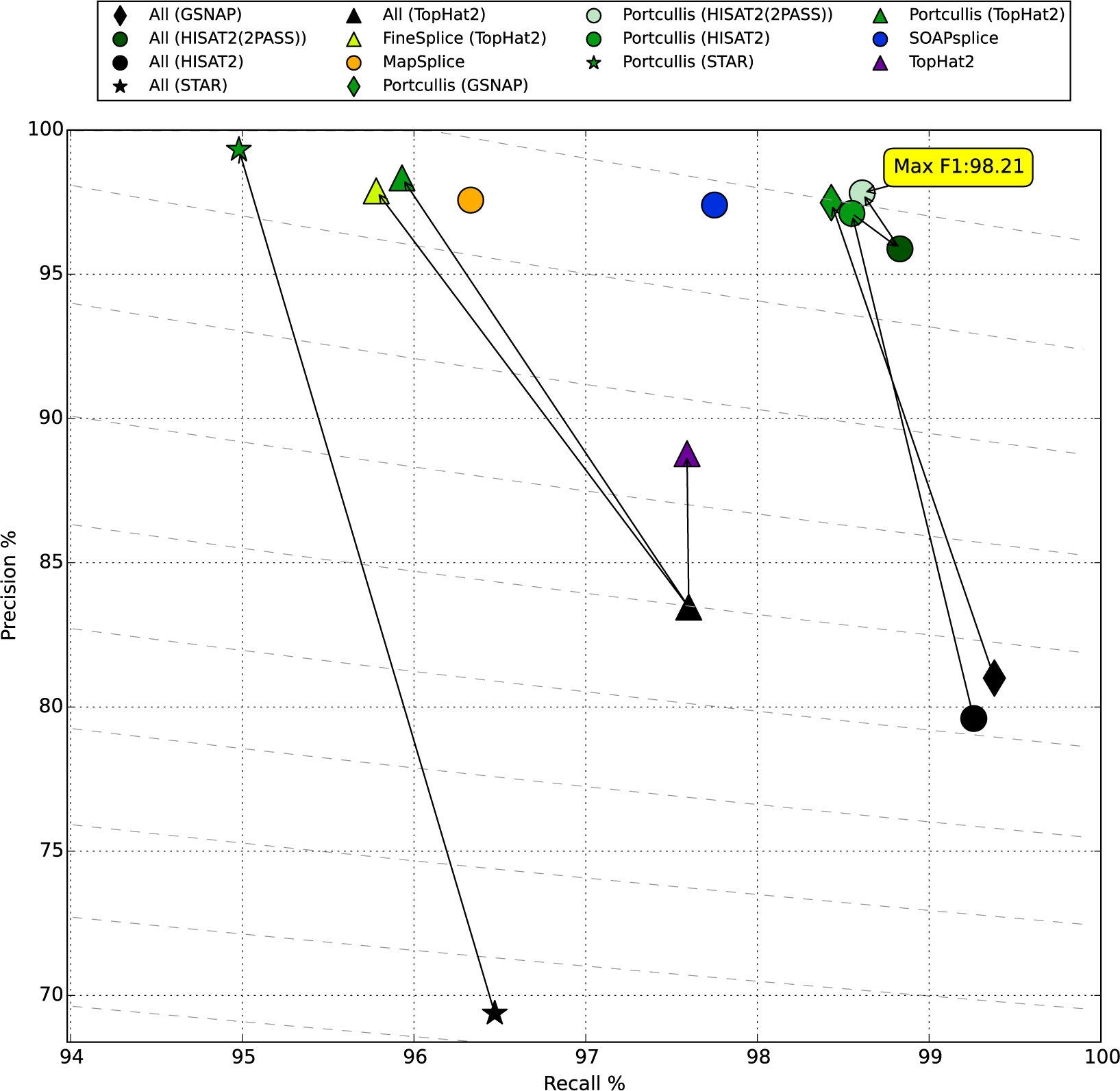
A scatter plot showing recall and precision results of all methods on our 101bp ~76 million simulated human read dataset. Diagonal lines represent actual *F*_1_ score gradients. Arrows show the effect of processing BAM files by downstream junction filtering tools such as Portcullis or FineSplice. The purple TopHat2 entry shows the effect of TopHat2’s own rule-based filtering on the BAM file.

**Fig. 5:**
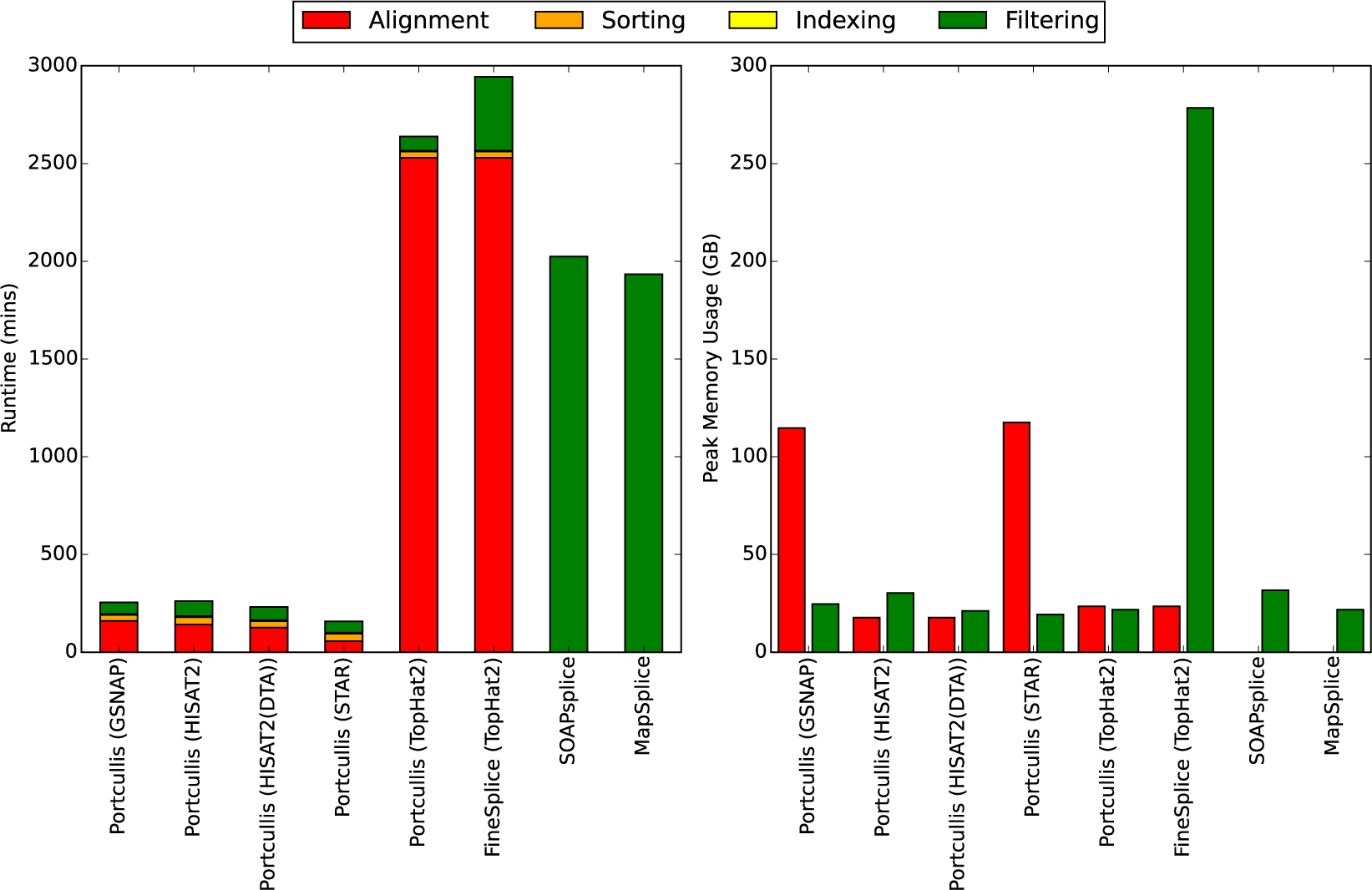
Runtimes and max memory usage of all methods on our 101bp ~76 million simulated human read dataset using 8 threads where appropriate. For FineSplice and Portcullis, times for alignment, sorting and indexing are factored into the results. For memory usage we consider alignment and filtering stages only.

An alternative approach to improving accuracy can be achieved by analysing the set of mapped split reads supporting each SJ and then applying some criteria to determine if that SJ is likely to be genuine or invalid. These criteria might involve using rules based on some pre-defined cutoff values, however, only modest gains can be achieved this way, see SI section 2 for details. More accurate results can be achieved through a more comprehensive analysis combining multiple metrics, which typically comes at the expense of time and computational resources. Some tools that do this include: FineSplice v0.2.2 (Gatto *et al.*, 2014), TrueSight v0.06 (Li *et al.*, 2013), MapSplice v2.2.1 (Wang *et al.*, 2010a) and SOAPsplice v1.10 (Huang *et al.*, 2011). These methods vary in their implementation and intended usage. MapSplice, TrueSight and SOAPsplice are standalone tools designed to be used as highly accurate RNAseq mapping tools. They process reads directly and produce alignments and lists of detected junctions. Finesplice is a post-alignment junction filtering tool, requiring a TopHat2 BAM file as input, which produces a list of filtered SJs.

However, as we will show, all these tools have specific disadvantages, and this prompted us to develop our own method, Portcullis. Portcullis has a similar architecture to Finesplice, consuming BAM files to produce a set of filtered SJs. In addition, portcullis analyses all SJs present in the input BAM, and can consume BAMs generated from any RNA-seq mapper, not just TopHat2. The Portcullis method is described in section 5.3.

We undertook an experiment to observe how these methods perform when varying sequencing depth, read length and species. The full results are presented in SI figures 6 to 8, although, we provide a recall / precision scatter plot in figure 4, which summarises many of the same points in a single plot. This plot shows how the methods compare when run on our 101bp simulated human dataset. The difference between the more accurate methods and the original RNA-seq mappers is stark, with precision scores not dropping below 97% as compared to precision lower than 85% in all cases for the mappers. Because Finesplice and Portcullis are dependent on the alignments produced by RNA-seq mapping tools, it is impossible for them to outperform their input in terms of recall, so their task is to discard invalid, but not genuine, junctions. Despite a small drop in recall, Portcullis significantly improves the precision of splice junction calls over all input methods producing higher overall *F*_1_ scores, and outperforms FineSplice on TopHat2 input. In addition, for this dataset when coupled with GSNAP or HISAT2 Portcullis produces the highest overall *F*_1_ scores of any method. For most methods increasing read length improves results, although while SOAPsplice has a comparable *F*_1_ score in this plot, SI figure 6 shows that increasing read length over 101bp decreases SOAPsplice’s accuracy drastically. Furthermore, we can improve *F*_1_ even more by running HISAT2 in a two pass configuration, feeding in portcullis predicted junctions in the second pass.

**Fig. 6:**
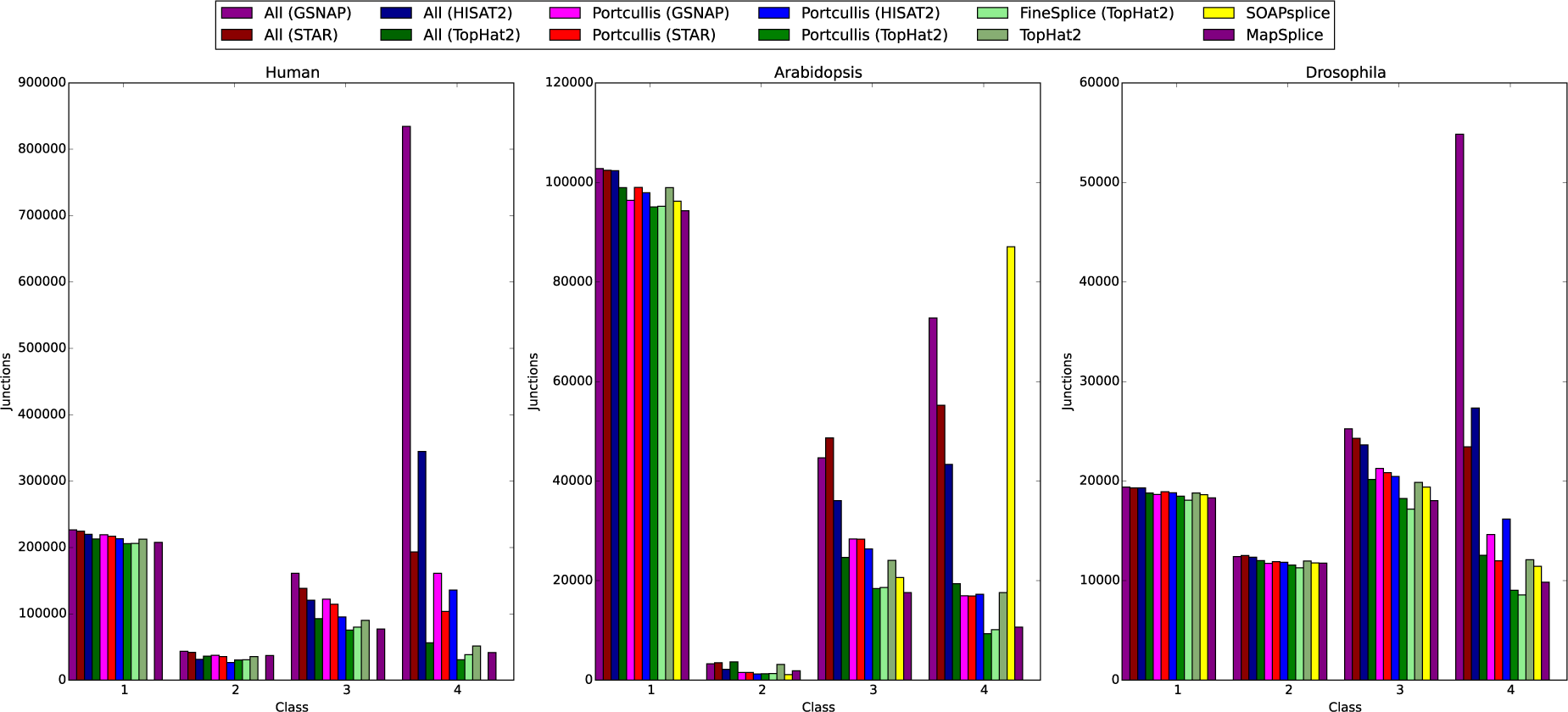
In this plot we check junctions found via each method against the reference annotation for Human (251bp reads), Arabidopsis (151bp reads) and Drosophila (101bp reads) respectively. The results are categorised into the following classes: 1) intron match; 2) both splice sites found; 3) one splice site found; 4) no splice sites found. SOAPsplice did not finish for the Human dataset and TrueSight could not be run on any.

The runtimes and max memory usage associated with methods that deliver high accuracy in figure 4 are shown in figure 5, with results across all datasets shown in SI figures 10 to 11. The results include time taken to align, sort and index the input reads where required, in addition to filtering the junctions. All methods were run with 8 threads where possible, and we would also like to point out that Portcullis’ memory usage can be lowered further by reducing the number of threads, as shown in SI figure 12.

This runtime analysis immediately highlights issues with several methods. First, TopHat2 suffers from long runtimes, which should be considered when running downstream filtering tools. This is especially problematic to FineSplice, which can only run on TopHat2 alignments. FineSplice also suffers from high memory usage, which grows quickly with the number of reads present in the dataset. High memory usage is also an issue with GSNAP, STAR and TrueSight. Indeed, we terminated the TrueSight job on our 101bp human dataset after a week of processing. Both SOAPsplice and MapSplice have relatively long runtimes compared to Portcullis coupled with GSNAP, STAR or HISAT2. Portcullis performs well, both in terms of accuracy and practicality, across varying sequencing depth, read length and species, particularly when coupled to HISAT2.

### 2.2 Analysis of real data

RNAseq simulations, while useful for benchmarking methods, do not give a full description of the complexity and noise inherent in real data. For example, our artificial datasets only simulated transcripts present in the reference annotations, which are heavily biased towards coding transcripts (Djebali *et al.*, 2012). Real data will contain many more non-coding elements and may contain other biological entities that have not been properly annotated yet.

We predict that portcullis is more useful on real data than on simulated, given that it appears to have a greater effect on noisier and deeper datasets. However, comparing the accuracy of methods on real data is more challenging for several reasons. Firstly, even for model organisms, the reference annotations are incomplete and RNA-seq experiments will likely contain some genuine novel junctions, which appear as false positives, meaning precision scores are unreliable. Furthermore, when comparing precision scores between methods it does not necessarily follow that higher precision is a better result, as this could be interpreted as the tool is worse at finding lower confidence junctions. Secondly, a single RNA-seq experiment is unlikely to cover all junctions found in the reference, meaning that it is impossible to achieve perfect recall, although typically we can compare recall scores between methods, as a higher number of correctly detected reference junctions suggests that the method is genuinely more sensitive.

To get a better feel for the validity of SJs on real data it is helpful to take a finer grained approach. For simulated data, SJs were considered as pairs, representing both ends of the intron and that SJ pair is considered genuine only if both SJs are found in the reference. Instead of considering just two classes (genuine and invalid) for this pairing it is possible to breakdown each intron into four distinct classes, each with decreasing likelihood of representing a genuine SJ (Li *et al.*, 2013):

1. introns matching annotated known introns (i.e. both splice sites match the same intron in the reference).
2. introns with donor and acceptor sites present in annotation but matching different introns
3. introns with only one annotated splice site (i.e. only one splice site found in the reference)
4. introns with two novel splice sites (i.e. both not found in the reference)

By classifying junctions this way we should be able to compare the precision of methods based on higher counts as the class number increases. To demonstrate this point, we extracted junctions from our simulated 76bp human dataset and compared them against the real human reference. The results shown in SI figure 13 indicate the majority of junctions occurring in class 1, with similar counts across methods. This is expected for simulated data as all correct junctions should only be found in class 1, with false positives seen by high counts in classes 2, 3 and 4.

To assess the performance of Portcullis on real data we used the same real datasets that provided error models for our simulated dataset and checked junctions predicted by various methods against the reference annotation for each species: Human, Arabidopsis and Drosophila. The results shown in figure 6 indicate that counts in class 1 have little variation between methods, across species, which implies that the methods that performed well on simulated data are not discarding many genuine junctions incorrectly. As expected, counts in class 4 have a high variation between methods across species. The methods identified as being most imprecise for our simulated data also have higher counts in this class relative to the more precise methods. This implies that much of the difference is likely explained by false positives. So the large drop between input mappers and Portcullis is reassuring and likely means that Portcullis’ predictions in this class are likelyto contain a higher proportion of genuine novel junctions than the input mappers. These plots also highlight again SOAPsplice’s problem handling long reads, as for the Human dataset (251bp) SOAPsplice failed completely, and with Arabidopsis (151bp) we can see a very high number of class 4 junctions. This is in contrast to the Drosophila dataset, which contains 101bp reads where SOAPsplice appears to perform well, in line with counts from Portcullis and MapSplice. We could not get TrueSight to run on any of these real datasets due to excessive memory runtime requirements.

## 3 USE CASE FOR NON-MODEL ORGANISMS

To demonstrate the use of Portcullis in a challenging real-life scenario, we created HISAT2 alignments for 6 Chinese Spring Wheat RNAseq samples (Clavijo *et al.*, 2017), producing a total 1,515,705,216 alignments of 251bp reads across all datasets. We aligned HISAT2 with and without HISAT2’s Downstream Transcript Assembly (DTA) mode, which is intended to reduce the number of false positive splice junctions in the aligned data. We then ran Portcullis on each set of aligned reads. Using Junctools (see section 5.3.4), we took the union of junctions across each set of samples, marking up the number of samples each junction occurs in, along with whether the junction can be found in the wheat reference annotation. Figure 7 shows that junctions found in all samples are also likely to be found in the reference annotation and have a high expression. Conversely, junctions detected in a single sample have a lower expression on average, meaning they are less likely to be incorporated into the reference annotation and are more likely to be false positives.

**Fig. 7:**
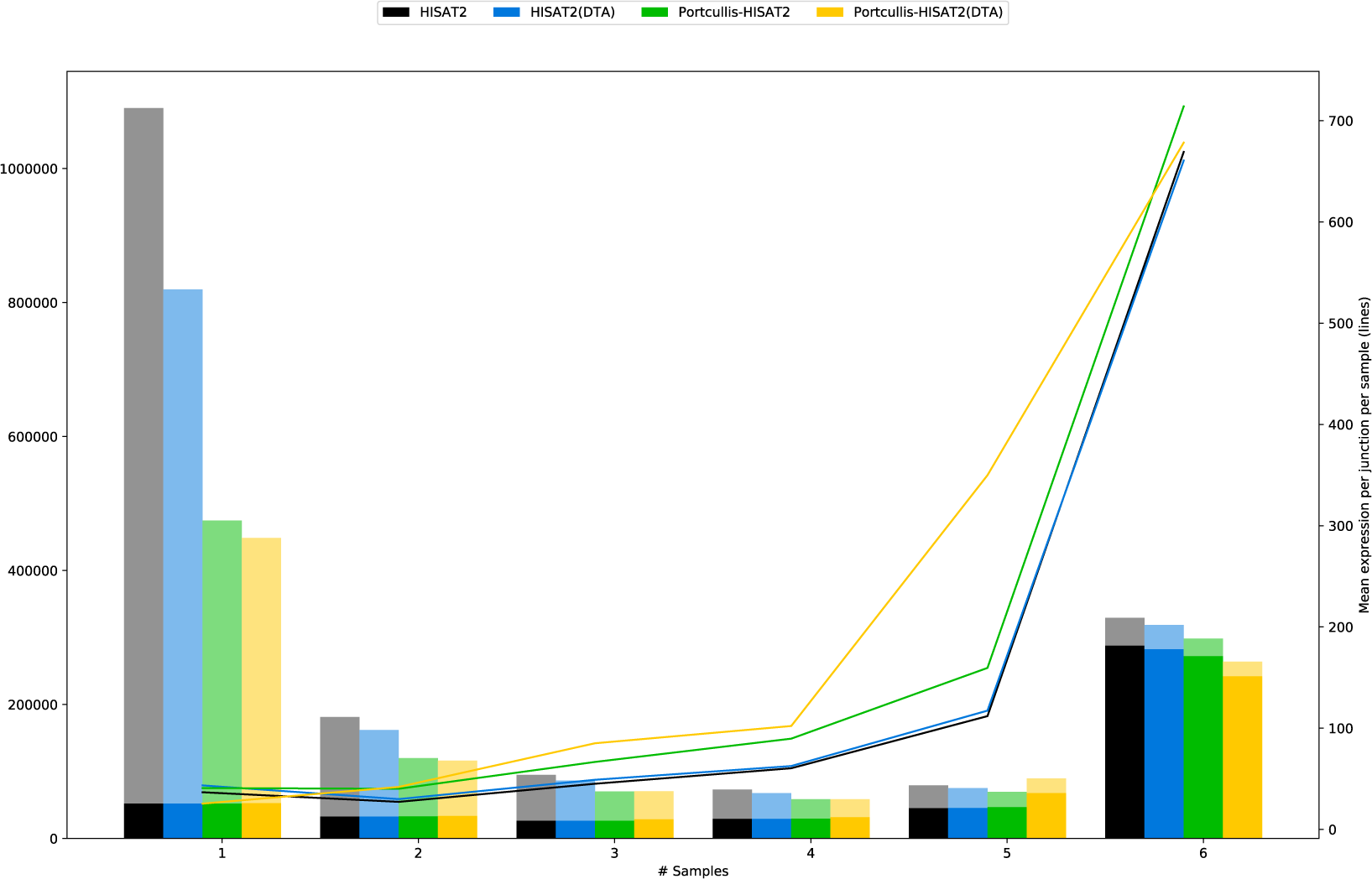
Junction counts that are supported by 1 through 6 samples for Wheat RNAseq data. Solid tint indicates that junctions were found in the reference annotation, paler tint indicates junctions were not found in reference. Junctions which occur in all 6 samples are more likely to be found in the reference. Average expression per junction per sample is shown by the lines, and indicates higher expression for junctions which occurs across all samples.

Reassuringly, the overall number of junctions for Portcullis in both modes is similar: 1, 091, 198 for HISAT2, and 1, 047, 335 for HISAT2(DTA), whereas the difference in the number of junctions found in the pre-filtered alignments was much larger: 1, 848, 057 and 1, 529, 999 respectively. Furthermore, the number of genuine junctions found in a single sample are consistent across all datasets, indicating that Portcullis is not falsely discarding many annotated, and therefore likely genuine, albeit lowly expressed, junctions.

Another interesting but subtle point that appears in Portcullis filtered junctions with HISAT2(DTA), is that we see a notable drop in those categorised in 6 samples, with a corresponding increase in the 5 sample category. This is caused by Portcullis filtering out a junction in 1 of the 6 samples.

In addition, to demonstrate Portcullis’ ability to handle large datasets, we merged the BAMs from all 6 samples, producing a single BAM containing alignments of ~755 million 251bp reads. Portcullis processed these in 400 minutes, using 4 threads, and requiring a peak of 59GB of RAM on a machine with 4 AMD Opteron(tm) 6134 processors. The full set of HISAT2 alignments consisted of 1, 845, 781 junctions with 1, 092, 390 junctions remaining after filtering.

## 4 DISCUSSION

In this manuscript we confirmed that recent versions of popular RNA-seq alignment tools still suffer from high numbers of false positive splice junctions in their output. Achieving the best results requires a more thorough and computationally intensive analysis of alignments around splice sites and interrogation of the genome at these loci. However, most other tools that achieve high accuracy, are either impractical or unreliable across different datasets and species. Portcullis is the only junction prediction tool that achieves these results, while scaling to the requirements of modern NGS workflows. We demonstrated that Portcullis is capable of analysing a wheat dataset comprising of ~755 million 251bp reads merged from 6 separate samples, on a fragmented version of the genome containing 735, 943 contigs. Portcullis ran to completion in 400 minutes using 4 threads with less than 60GB of RAM, making it feasible to process extremely large, complex datasets with readily available hardware.

Apart from accuracy and runtime performance, we see one of the strengths of Portcullis is its flexibility and ease of use. Especially notable is the fact that it completely decouples junction filtering from RNAseq mapping, enabling the user to select the mapper of their choice, should more attractive mapping options be available in the future.

In addition, Portcullis’ supplementary toolkit ’Junctools’, ensures that Portcullis’ output is easy to incorporate into workflows that use other tools, reducing the amount of custom scripting required by a bioinformatician. Junctools, also makes it straightforward to integrate Portcullis into a two-pass alignment approach, whereby portcullis junctions are converted to a format suitable for a particular aligner via Junctools, then used as as a guide to produce more accurate alignments. In particular, we see that coupling HISAT2 with Portcullis in two-pass mode, delivers high accuracy and is still within a reasonable time-frame and acceptable memory usage.

Accurate SJ prediction allows us to get richer and more useful information from downstream tasks such as alternative splicing analysis. Each missed SJ reduces the richness of the transcript model and each false positive can lead to incorrect donors, acceptors and cassettes. This is particularly important in non-model organisms with incomplete annotations. In addition, transcript assemblies could be improved either by filtering transcripts containing unsupported SJs, or by producing more accurate input reads via the two-pass approach mentioned previously. Portcullis junctions can be used to provide additional hints to gene prediction tools to help select between sets of alternative isoforms. This way we envisage Portcullis assisting the production of both richer and more precise genome annotations for Eukaryotes.

## 5 MATERIALS AND METHODS

### 5.1 Simulation of RNA-seq data

To compare performance between junction filtering tools we created several simulated RNA-seq datasets based on three known model organisms (Human, Drosophila and Arabidopsis), with error and expression profiles derived from real datasets using SPANKIsim (Sturgill *et al.*, 2013). This produces reads derived from a known region in the reference transcriptome, along with the perfect alignments of those reads. From the alignments it is possible to unambiguously derive the true set of junctions for the given dataset, providing a platform from which RNA-seq mappers and SJ filtering tools can be benchmarked. The complete pipeline used to generate the simulated reads is described in SI section 1.2. Basic statistics for the datasets are described in table 1.

**Table 1.** Properties of core simulated datasets for each species

| Properties of simulated dataset | Arabidopsis | Drosophila | Human |
| --- | --- | --- | --- |
| Reference annotation | TAIR10 | DMEL78 | HG38 |
| Original accession | PRJEB7093 | SRA009364 | PRJEB4208 |
| Millions of reads (in original) | 93 | 47 | 97 |
| Mean quality (in original) | 37 | 37 | 39 |
| Depth fractions | 10%,50%,100% | 10%,50%,100% | 10%,50%,100%,200% |
| Millions of paired reads | 9,47,93 | 5,24,47 | 7,38,76,152 |
| Read lengths @ 100% depth (bp) | 76,101 | 76 | 76,101,151,201 |
| # splice junctions @ max readlen and 100% depth | 90,190 | 29,275 | 139,403 |
| % of SJ's in ref | 71% | 51% | 42% |
| # transcripts @ max readlen and 100% depth | 19723 | 9376 | 19853 |
| % of transcripts in ref | 47% | 32% | 12% |

### 5.2 Performance Evaluation

For most of the experiments described in this paper, our truth set consists of a subset of genuine junctions taken from a reference transcriptome. As it is impractical to derive a comprehensive set of false junctions (every combination of start and stop sites in the genome that are not genuine junctions) we use performance metrics more commonly associated with information retrieval: recall (equation (1)), precision (equation (2)) and *F*_1_ measure (equation (3)). The equations use counts of: true positives (*TP*); false negatives (*FN*); and false positives (*FP*). Note that there are no true negatives (*TN*) for reasons mentioned previously. Recall represents the fraction of true junctions retrieved, otherwise known as sensitivity or the true positive rate. Precision represents how often when the system makes a call it is correct, otherwise known as the positive predictive value. *F*_1_ is the harmonic mean of recall and precision and a convenient proxy for the overall accuracy of the system.

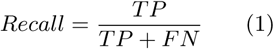

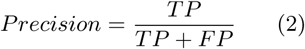

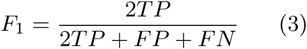

## 5.3 Portcullis

The pipeline for Portcullis, shown in figure 8, describes the flow of data from sequenced RNA reads in FastQ format through to a set of filtered junctions provided by Portcullis. RNAseq files must first be mapped with a suitable RNAseq mapper and converted into BAM format. Portcullis takes in one or more BAM files and a genome in FastA format as input. The first stage in Portcullis ensures all input data is prepared in a way suitable for downstream processing. This includes, if multiple BAM files are provided merging them into a single file, then ensuring that the BAM is coordinate sorted, and ensuring that both the BAM and genome file are indexed. All distinct junctions are extracted from the BAM file and put into a set and analysed. This process is described in detail in section 5.3.1. The full set of junctions is then put into the Portcullis filter tool, which is described in more detail in section 5.3.3. The output is both the full set and the filtered set of junctions in both a descriptive tabular format and an exon-based bed format, both of which can be used in downstream analyses.

**Fig. 8:**
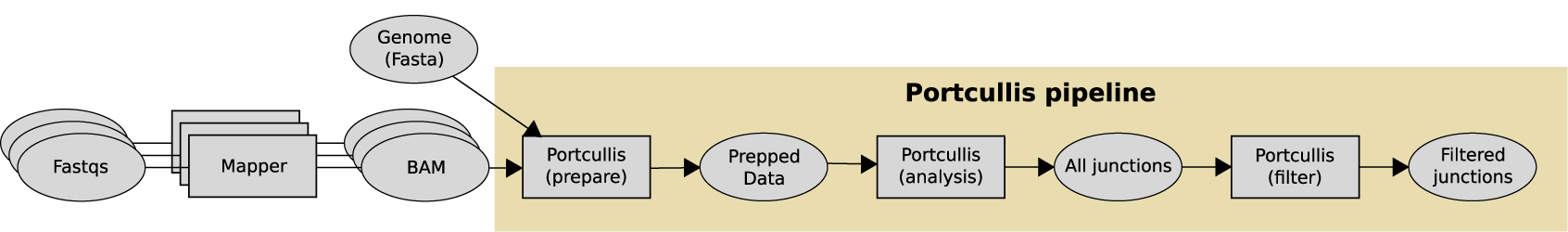
A high level view of the Portcullis pipeline. Input to Portcullis is a genome in fasta format and one or more BAM files created by an upstream RNAseq mapping tool. The first stage ensures the alignments are correctly merged, sorted and indexed, then all junctions found in the input are analysed and output to disk. Next, the full set of junctions are filtered to remove likely false positives and are also output to disk. The user can either choose to run the full pipeline in one go, or each stage separately.

### 5.3.1 Junction Analysis

The first step to analyse junctions present in the BAM is to identify split reads containing reference skipping cigar operations (’N’), then to collapsse these reference skipping regions into a distinct set of junctions defined by their location (target sequence and start site) and size. Portcullis makes observations about each potential distinct junction based on the RNA-seq data and the genomic features at those loci. A definitive, and up to date list of features is described with the software’s documentation, although we describe a few of the more interesting features here.

First, we provide a number of ways to quantify each splice junction. The most obvious metric is a raw count of the number of split reads supporting the junction. Another is to count how many of those split reads contain only a single intron (uniquely split reads - USRs), and how many are uniquely mapped (typically having a high MapQ score) (uniquely mapped split reads - UMRs). We consider a split read to be ’reliable’ if it maps uniquely, and is properly paired (it’s paired read is upstream or downstream on the expected strand of the current chromosome and within an expected distance). In addition, we find that that junctions possessing high ratios of reliable split reads to the total number of raw split reads seems to be a useful indicator of junction quality.

Assuming random sampling of sequenced reads, it should be expected that the start sites of split reads should be uniformally distributed across the upstream anchor of the junction. This notion is captured in the Shannon entropy score (Wang *et al.*, 2010a). Junctions with high number of split reads can therefore have a low entropy score if those reads start at a small number of sites and are therefore less likely to be genuine. Similarly junctions with a moderate number of supporting reads can have high entropy (and therefore more likely to be genuine) if many of them have distinct start sites. This concept is extended further to show the deviation from expected to observed read counts at each anchor position (Gatto *et al.*, 2014), providing a more detailed picture across the split read overhangs up and downstream of the intron. Typically we expect a gradual reduction of coverage across the length of the junction (up to half the read length), where we see sharp deviations from this, the chance of an invalid junction increases.

Another frequently used approach is to calculate the maximal split read overhang, an method used in both TopHat2 and STAR for filtering junctions. The best score possible for a given junction is the maximum split read length divided by 2. A more sophisticated version of this concept includes modifying this score by penalising alignments containing mismatches. This approach is called the maximum of the minimal match on either side of the splice junction (MaxMMES) (Wang *et al.*, 2010b).

In terms of genomic information, we consider the composition of the two-base donor and acceptor sites, which are in most, but not all, cases the conform to the same canonical, or semi-canonical, pattern (Burset *et al.*, 2000) associated with the minor spliceosome. We also look at hamming distances between the left anchor and right side of the intron, as well as the left intron and right anchor. This provides an indication of whether the splice sites are embedded in a repeat region (Sturgill *et al.*, 2013) and therefore unlikely to be genuine. This idea is illustrated in figure 9.

**Fig. 9:**
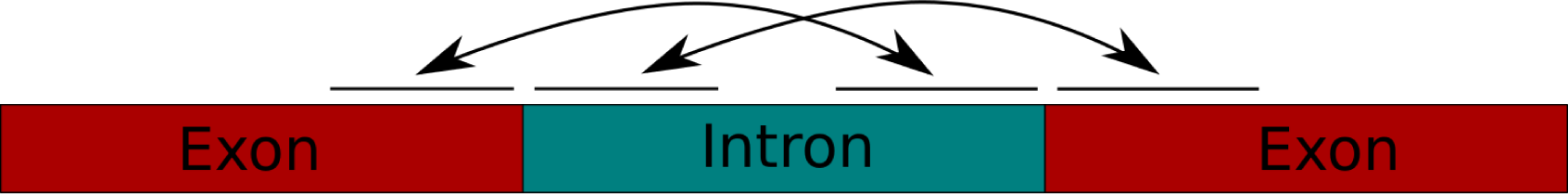
Calculating the hamming distinct of both the rightmost region of the left anchor to the rightmost region of the intron, and the leftmost region of the intron to the left most region of the right anchor can give an indication of whether the splice site may have been incorrectly triggered by a repeat region in the genome.

### 5.3.2 Feature extraction and analysis

It’s possible to collect a highly precise subset of SJs across different species and datasets through aggressive rule-based filtering section 2. Similarly, we can collect a high confidence subset of invalid junctions through an alternative set of rules. By using information present in both subsets it is possible to extract additional features which can be retrospectively applied to all junctions in the dataset. Features derived this way are described here:

- Intron score (Li *et al.*, 2013). SJs with excessively long introns are likely to be incorrect, however, the average intron sizes of splice junctions vary dramatically between species (Zhu *et al.*, 2009). Therefore, in order to derive useful information from the intron size, we need to compare the intron size against those found in our initial positive set. Introns greater than that found in the 95th percentile of our positive set are assigned a score.
- Splicing signal (Li *et al.*, 2013). Many *ab initio* gene prediction tools use markov chains to predict splicing signals derived from the genome alone (Stanke and Waack, 2003). The idea is to use the high-confidence SJs to create a markov chains modelling the donor and acceptor sites. From there we can create a markov model that defines a splicing signal score for each SJ in our full set.

To show that these additional features provide additional discriminative power we ran PCA and tSNE on HISAT2 alignments of the 201bp simulated human read dataset. Both methods were run before and after the additional extracted features are added. The results are shown in the first two columns of figure 10. The left column (showing results without extracted features) has more overlap between genuine and invalid junctions than the middle column (with extracted features).

**Fig. 10:**
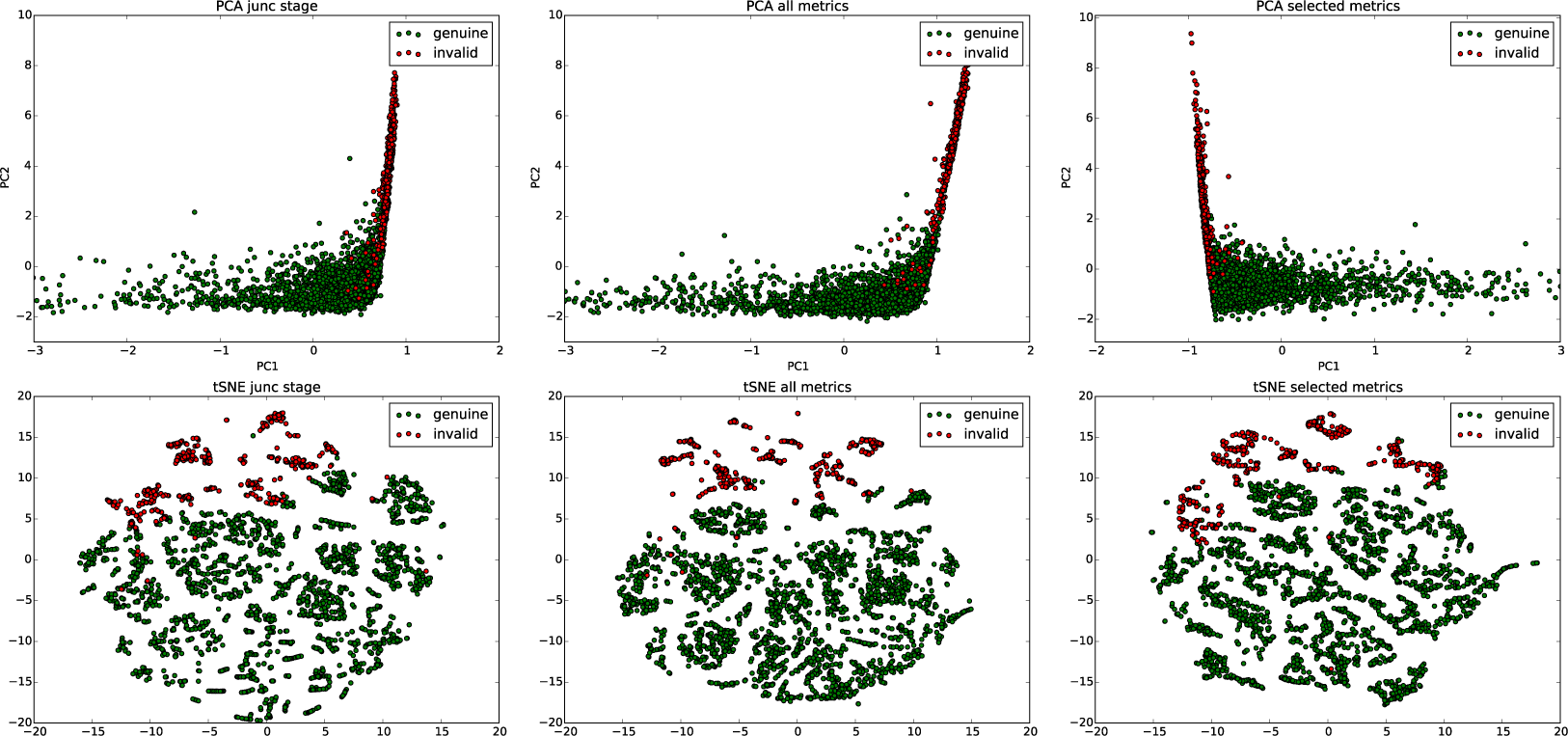
The plots show separation of valid and invalid junctions in HISAT2 alignments of simulated 201bp reads from the human dataset. The top row shows PCA plots and the bottom row shows tSNE plots. The left column shows the all metrics prior to the filtering stage without additional extracted features. The middle column shows the full set of metrics including extracted features. And the right hand column shows the subset of features used in our classifier.

However, while, in general, supplying additional features improves predictive performance, sometimes additional features increases the chance of a classifier overfitting to the data when training (Hira and Gillies, 2015). A classifier can often generalise better and run significantly faster with a smaller feature set. We discarded features that we determined to not be useful for classification purposes by running our classifier through permutations of features and removing those, which had no, or a negative, change in accuracy across all our simulated datasets, and repeated this process until no further gains we apparent. The features that passed our feature selection step are listed below:

- Reliable reads
- Reliable to raw read ratio
- MaxMMES
- Mean mismatches per read
- Intron score
- Min hamming score
- Position weight matrix
- Splicing signal
- Deviation of expected to observed read counts at each anchor position

To show that discriminative power wasn’t diminished we show the PCA and tSNE plots for this data in the right column in figure 10.

### 5.3.3 Filtering junctions via a self-training classifier

The Portcullis filtering pipeline, shown in figure 11, bears some similarities with the approaches used by TrueSight (Li *et al.*, 2013) and FineSplice (Gatto *et al.*, 2014) in that it combines information derived from both the genome and RNA-seq mapping features and an initial positive and negative training set to train a model that can then be applied to assign a score to every junction in the input dataset. Because this approach trains on the same data its making predictions on, it helps to make the filtering approach agnostic to read length, read quality, sequencing depth, species and mapping tool.

**Fig. 11:**
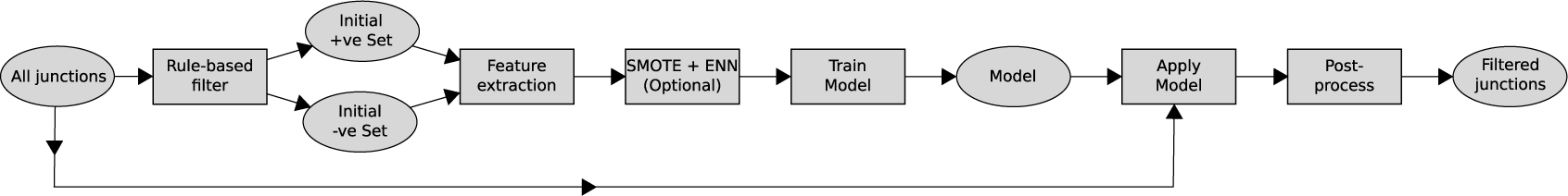
An exploded view of the Portcullis filtering stage. Input is a set of junctions to filter in tab format. This pipeline first creates a model from a high confidence set of likely genuine and likely false junctions. The model is then applied to the full set of junctions and output in tab and bed format.

Prior to training, the initial sets are then balanced using SMOTE (Chawla *et al.*, 2002), a synthetic oversampling technique, in order to partially compensate for biases that can be introduced that favour the larger set. The balanced training set is then used to train a random forest (Breiman, 2001) using 100 trees (see SI section 3 for justification), which is then applied to the full set of junctions in order to assign a probability score. Typically, values *>*= 0.5 are used to define genuine junctions and *<* 0.5 as invalid, although we leave the actual threshold used as an option for the user in case they would like to prioritise recall or precision.

### 5.3.4 Junctools

Portcullis comes bundled with a supplementary toolkit called *Junctools*, which provides the user with a number of features for manipulating and analysing junction files in many commonly used formats.

*Convert* - Junctools can convert between commonly used junction files, such as BED and GFF, or between these and input to guide several popular RNA-seq mappers so they can be easily driven in two-pass mode without requiring the user to do their own scripting.

*Compare* - Junctools can compare junction files against each other, which can be useful to see how many junctions from Portcullis are present in a given reference.

*Set operations* - In addition, Junctools supports set operations between junction files, which can be useful for merging results between several Portcullis runs, or to separate junctions not found in a reference annotation for example.

*GTF* - Finally Junctools offers the user the ability to filter out or markup transcripts in GTF files, that do not contain junctions found in a separate junctions file. This allows the user to filter out transcripts that do not have junctions that are supported by Portcullis for example.

## ACKNOWLEDGEMENTS

This research was supported in part by the NBIP Computing infrastructure for Science (CiS) group.

## Supplementary Information

Supplementary Information (SI) is available online. In addition, the software documentation is available online at: http://portcullis.readthedocs.io/en/latest/

